# Molecular Characterization, Phylogenetic and Variation Analyzes of SARS-CoV-2 strains in Turkey

**DOI:** 10.1101/2020.09.11.293183

**Authors:** Karamese Murat, Ozgur Didem, Tutuncu Emin Ediz

**Author notes:** **Corresponding Author**, Murat KARAMESE, Kafkas University, Medical Faculty, Department of Microbiology, 36100, Kars, Turkey.

## Abstract

**Introduction:** We present the sequence analysis for 47 complete genomes for SARS-CoV-2 isolates on Turkish patients. To identify their genetic similarity, phylogenetic analysis was performed by comparing the worldwide SARS-CoV-2 sequences, selected from GISAID, to the complete genomes from Turkish isolates. In addition, we focused on the variation analysis to show the mutations on SARS-CoV-2 genomes.

**Methods:** Illumina MiSeq platform was used for sequencing the libraries. The raw reads were aligned to the known SARS-CoV-2 genome (GenBank: MN908947.3) using the Burrows-Wheeler aligner (v.0.7.1). The phylogenetic tree was constructer using Phylip v.3.6 with Neighbor-Joining and composite likelihood method. The variants were detected by using Genome Analysis Toolkit-HaplotypeCaller v.3.8.0 and were inspected on GenomeBrowse v2.1.2.

**Results:** All viral genome sequences of our isolates was located in lineage B under the different clusters such as B.1 (n=3), B.1.1 (n=28), and B.1.9 (n=16). According to the GISAID nomenclature, all our complete genomes were placed in G, GR and GH clades. Five hundred forty-nine total and 53 unique variants were detected. All 47 genomes exhibited different kinds of variants. The distinct variants consist of 274 missense, 225 synonymous, and 50 non-coding alleles.

**Conclusion:** The results indicated that the SARS-CoV-2 sequences of our isolates have great similarity with all Turkish and European sequences. Further studies should be performed for better comparison of strains, after more complete genome sequences will be released. We also believe that collecting and sharing any data about SARS-CoV-2 virus and COVID-19 will be effective and may help the related studies.

## Introduction

A new virus-associated disease named as coronavirus disease-2019 (COVID-19) by Severe Acute Respiratory Syndrome Coronavirus-2 (SARS-CoV-2) was firstly declared in late December of 2019 in Wuhan, China (1). Right now, there have been more than 16 million confirmed infections and over 645.000 deaths reported worldwide (2). The etiological agent of this disease, SARS□CoV□2 virus has a ssRNA (single □ stranded) genome and its length is approximately 29,890 base-pairs (bps) according to the data obtained from NCBI GenBank (GenBank NC_045512.2) (3). After the confirmation of human-to-human transmission and extensive global spread, World Health Organization (WHO) declared COVID-19 as a pandemic on the 11^th^ of March, 2020 (4).

To understand the transmission patterns and evolution of SARS-CoV-2 virus is crucial and necessary for creating better drug and vaccine designs for disease control and prevention (4–6). The analysis of genetic sequence data from a pathogen is known as an important tool in infectious disease epidemiology (7, 8). There are nearly 73.000 SARS-CoV-2 complete genome sequences available on the database of Global Initiative on Sharing All Influenza Data (GISAID) (9). The availability of these genomic data has been helping hundreds of researchers to analyze the genomic diversity of SARS-CoV-2 virus. Firstly, Rambaut et al. (8) have defined the lineage A and B (both of them includes sublineages such as A.1, B.1 or A.1.1) to help for tracking and understanding the patterns and determinants of the global spread of SARS-CoV-2. Then, at 4^th^ of July, 2020, GISAID published the new clade and lineage nomenclature aids in genomic epidemiology studies of SARS-CoV-2 viruses. They developed a nomenclature system based on marker mutations in phylogenetic groups and named as S, L, V, G, GH and GR (10).

In this study, we performed the NGS analysis for 47 complete genomes for SARS-CoV-2 isolates on Turkish patients. To identify their genetic similarity, phylogenetic analysis was performed by comparing the worldwide SARS-CoV-2 sequences, selected from GISAID, to the complete genomes from Turkish isolates. In addition, we focused on the variation analysis to show the mutations on SARS-CoV-2 genomes.

## Methods

This study was approved by both the Republic of Turkey Ministry of Health COVID-19 Scientific Research Evaluation Commission (Approval date: 02/05/2020; number: 2020-05-02T16_13_50) and the Local Ethics Committee of Kafkas University Faculty of Medicine (Approval date: 06/05/2020 number: 80576354-050-99/130).

Of the patients who tested positive for SARS-CoV-2, the 63 samples with the highest viral loads (assessed by Real time PCR targeting RdRp gene) were selected for Next-Generation Sequencing (NGS)-based assays. Fortyseven of those samples were used for complete genome sequencing and variation analyzes.

The library preparation was performed using CleanPlex® SARS-CoV-2 Panel (Paragon Genomics Inc., Iowa, USA) with isolated RNA samples following the manufacturer instructions. Then, Illumina MiSeq (Illumina, USA) platform with a 2×150-cycle kit was used for sequencing the libraries. The quality of the raw data was examined by FastQC v.0.11.5, and low-quality bases and primers were trimmed using Trimmomatic (version 0.32). The raw reads were aligned to the known SARS-CoV-2 genome (GenBank Accession: MN908947.3) using the Burrows-Wheeler aligner (version 0.7.1). The consensus sequences were aligned via multiple sequence alignment using MAFFT v7.450 tool. The phylogenetic tree was constructer using Phylip v.3.6 with Neighbor-Joining and composite likelihood method and 100 bootstrap replications. For phylogenetic analysis of our data, GISAID database was used to collect SARS-CoV-2 complete genomes of different patients all around the world (i.e. Belgium, Bosnian-Herzegovina, Czech Republic, Croatia, Cyprus, Finland, France, Germany, Hungary, Italy, Poland, Portugal, Spain, Russia, Australia, USA, Mexico, Costa Rica, Canada, Austria, Thailand, China, Kuwait, Senegal, Egypt, Japan, Chile, Nigeria, Algeria, Brazil, Ecuador, Mongolia, Colombia, India, Indonesia, Malaysia, Bangladesh, Kazakhstan, and Turkey). Only complete genomes (28,000–30,000 bps) were analyzed. The variants were detected by using Genome Analysis Toolkit-HaplotypeCaller (GATK) v.3.8.0 and were inspected on GenomeBrowse v2.1.2 (GoldenHelix). The variants with low quality and a low variant fraction (%<60) were removed. The filtered variants and reference SARS-CoV-2 genome were used to generate the consensus sequence using bcftools v1.9.

## Results

A database of 68 complete genome of SARS-CoV-2 strains belonging to different countries randomly selected from the GISAID database were compared to our isolations. The reference SARS-CoV-2 strain, which was rooted from Wuhan (NCBI GenBank, NC_045512) was also selected for comparison. A total of 115 SARS-CoV-2 genomes were placed in the phylogenetic tree. The maximum likelihood phylogenetic tree in Figure 1 and Figure 2 shows a main lineage including several sublineages. All viral genome sequences of our isolates was located in lineage B under the different clusters such as B.1 (n=3), B.1.1 (n=28), and B.1.9 (n=16). According to the GISAID nomenclature, all our complete genomes were placed in G, GR and GH clades (Figure 1, Figure 2 and Table 1).

**Figure 1.**
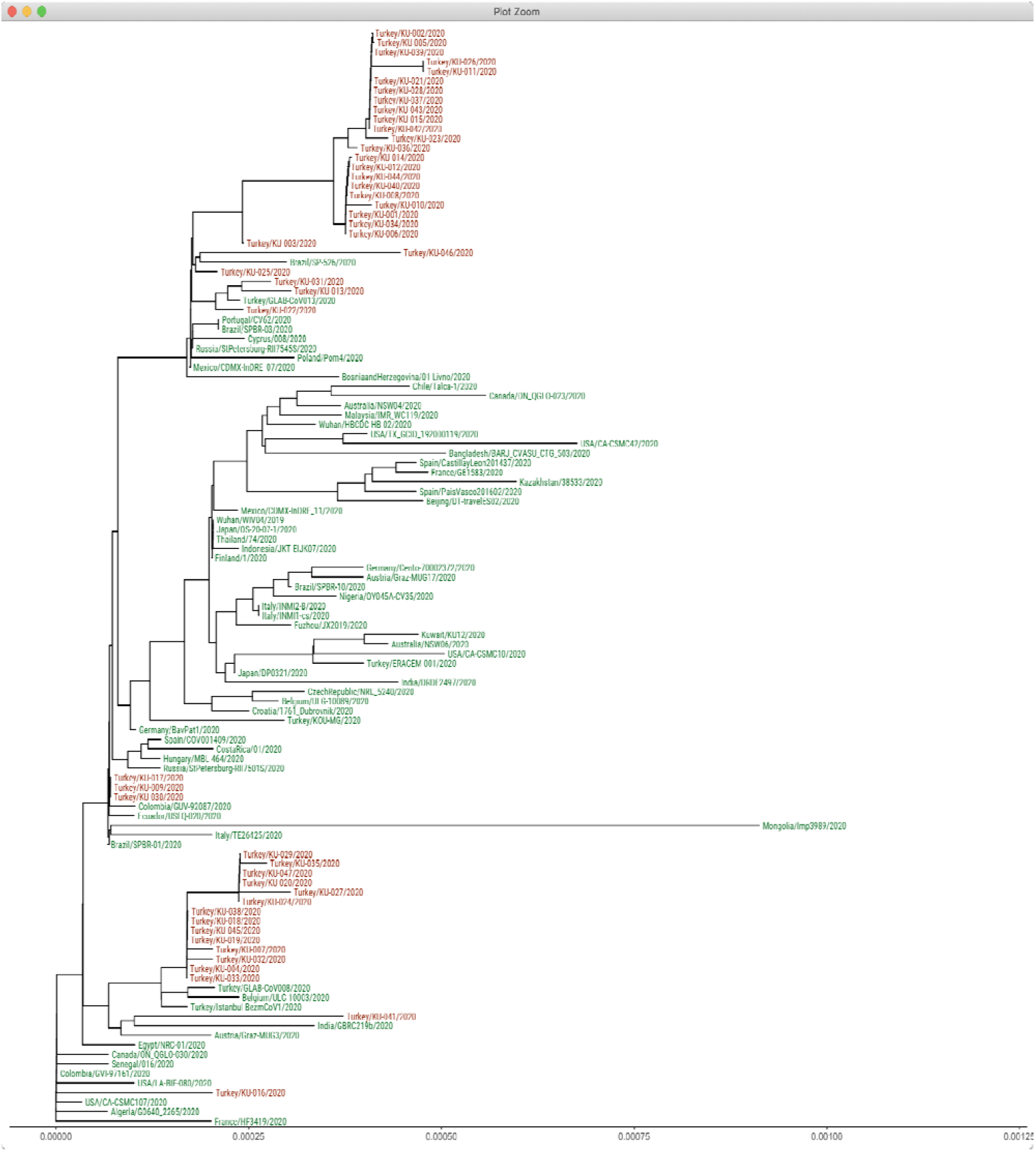
Global diversity of circulating SARS-CoV-2 strains including at least one strain in different countries. Our viruses are shown in red font color.

**Figure 2.**
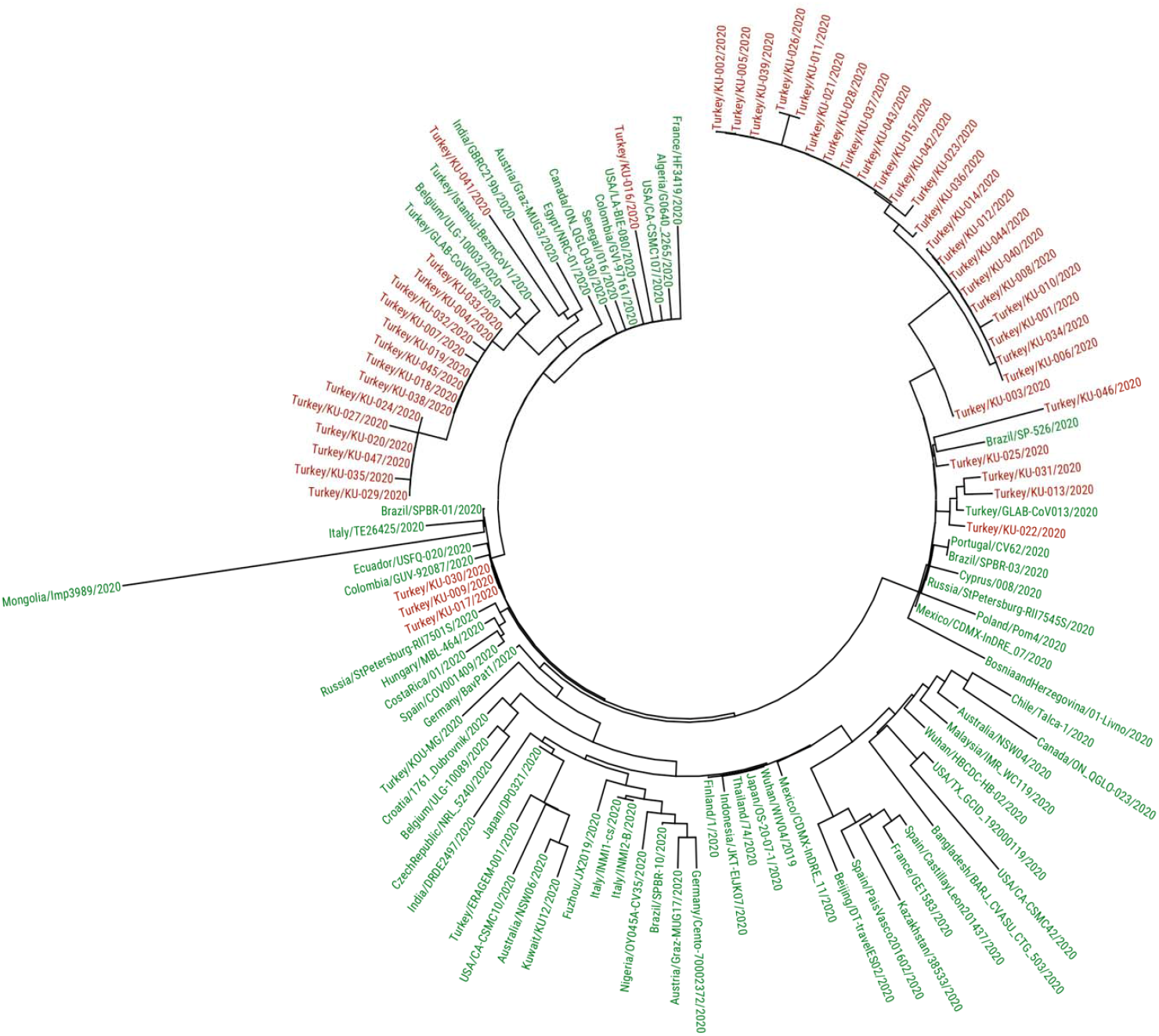
Sub-tree showing the SARS-CoV-2 strains containing also at least one strain in different countries. Our viruses are shown in red font color.

**Table 1:** The accession numbers obtained from GenBank and GISAID database ad GISAID clades of our isolates

| GenBank Name | GenBank Accession Number | GISAID Name | GISAID Accession Number | GISAID Clade |
| --- | --- | --- | --- | --- |
| Kafkas-SARSCoV2-001 | MT786327 | hCoV-19/Turkey/KU-001/2020 | EPI_ISL_495411 | B.1.1 (GR) |
| Kafkas-SARSCoV2-002 | MT786861 | hCoV-19/Turkey/KU-002/2020 | EPI_ISL_495412 | B.1.1 (GR) |
| Kafkas-SARSCoV2-003 | MT786865 | hCoV-19/Turkey/KU-003/2020 | EPI_ISL_495413 | B.1.1 (GR) |
| Kafkas-SARSCoV2-004 | MT786329 | hCoV-19/Turkey/KU-004/2020 | EPI_ISL_495414 | B.1.9 (GH) |
| Kafkas-SARSCoV2-005 | MT786797 | hCoV-19/Turkey/KU-005/2020 | EPI_ISL_495415 | B.1.1 (GR) |
| Kafkas-SARSCoV2-006 | MT786331 | hCoV-19/Turkey/KU-006/2020 | EPI_ISL_495416 | B.1.1 (GR) |
| Kafkas-SARSCoV2-007 | MT786334 | hCoV-19/Turkey/KU-007/2020 | EPI_ISL_495417 | B.1.9 (GH) |
| Kafkas-SARSCoV2-008 | MT786859 | hCoV-19/Turkey/KU-008/2020 | EPI_ISL_495418 | B.1.1 (GR) |
| Kafkas-SARSCoV2-009 | MT786860 | hCoV-19/Turkey/KU-009/2020 | EPI_ISL_495419 | B.1 (G) |
| Kafkas-SARSCoV2-010 | MT786335 | hCoV-19/Turkey/KU-010/2020 | EPI_ISL_495420 | B.1.1 (GR) |
| Kafkas-SARSCoV2-011 | MT786868 | hCoV-19/Turkey/KU-011/2020 | EPI_ISL_495421 | B.1.1 (GR) |
| Kafkas-SARSCoV2-012 | MT787647 | hCoV-19/Turkey/KU-012/2020 | EPI_ISL_495422 | B.1.1 (GR) |
| Kafkas-SARSCoV2-013 | MT787564 | hCoV-19/Turkey/KU-013/2020 | EPI_ISL_495423 | B.1.1 (GR) |
| Kafkas-SARSCoV2-014 | MT786866 | hCoV-19/Turkey/KU-014/2020 | EPI_ISL_495424 | B.1.1 (GR) |
| Kafkas-SARSCoV2-015 | MT787648 | hCoV-19/Turkey/KU-015/2020 | EPI_ISL_495425 | B.1.1 (GR) |
| Kafkas-SARSCoV2-016 | MT787650 | hCoV-19/Turkey/KU-016/2020 | EPI_ISL_495426 | B.1 (GH) |
| Kafkas-SARSCoV2-017 | MT787578 | hCoV-19/Turkey/KU-017/2020 | EPI_ISL_495427 | B.1 (G) |
| Kafkas-SARSCoV2-018 | MT787645 | hCoV-19/Turkey/KU-018/2020 | EPI_ISL_495428 | B.1.9 (GH) |
| Kafkas-SARSCoV2-019 | MT787577 | hCoV-19/Turkey/KU-019/2020 | EPI_ISL_495429 | B.1.9 (GH) |
| Kafkas-SARSCoV2-020 | MT787646 | hCoV-19/Turkey/KU-020/2020 | EPI_ISL_495430 | B.1.9 (GH) |
| Kafkas-SARSCoV2-021 | MT787582 | hCoV-19/Turkey/KU-021/2020 | EPI_ISL_495431 | B.1.1 (GR) |
| Kafkas-SARSCoV2-022 | MT787581 | hCoV-19/Turkey/KU-022/2020 | EPI_ISL_495432 | B.1.1 (GR) |
| Kafkas-SARSCoV2-023 | MT786336 | hCoV-19/Turkey/KU-023/2020 | EPI_ISL_495433 | B.1.1 (GR) |
| Kafkas-SARSCoV2-024 | MT786337 | hCoV-19/Turkey/KU-024/2020 | EPI_ISL_495434 | B.1.9 (GH) |
| Kafkas-SARSCoV2-025 | MT787580 | hCoV-19/Turkey/KU-025/2020 | EPI_ISL_495435 | B.1.1 (GR) |
| Kafkas-SARSCoV2-026 | MT789689 | hCoV-19/Turkey/KU-026/2020 | EPI_ISL_495436 | B.1.1 (GR) |
| Kafkas-SARSCoV2-027 | MT787500 | hCoV-19/Turkey/KU-027/2020 | EPI_ISL_495437 | B.1.9 (GH) |
| Kafkas-SARSCoV2-028 | MT787749 | hCoV-19/Turkey/KU-028/2020 | EPI_ISL_495438 | B.1.1 (GR) |
| Kafkas-SARSCoV2-029 | MT787745 | hCoV-19/Turkey/KU-029/2020 | EPI_ISL_495439 | B.1.9 (GH) |
| Kafkas-SARSCoV2-030 | MT787748 | hCoV-19/Turkey/KU-030/2020 | EPI_ISL_495440 | B.1 (G) |
| Kafkas-SARSCoV2-031 | MT787742 | hCoV-19/Turkey/KU-031/2020 | EPI_ISL_495441 | B.1.1 (GR) |
| Kafkas-SARSCoV2-032 | MT787486 | hCoV-19/Turkey/KU-032/2020 | EPI_ISL_495442 | B.1.9 (GH) |
| Kafkas-SARSCoV2-033 | MT787553 | hCoV-19/Turkey/KU-033/2020 | EPI_ISL_495443 | B.1.9 (GH) |
| Kafkas-SARSCoV2-034 | MT787473 | hCoV-19/Turkey/KU-034/2020 | EPI_ISL_495444 | B.1.1 (GR) |
| Kafkas-SARSCoV2-035 | MT787747 | hCoV-19/Turkey/KU-035/2020 | EPI_ISL_495445 | B.1.9 (GH) |
| Kafkas-SARSCoV2-036 | MT787743 | hCoV-19/Turkey/KU-036/2020 | EPI_ISL_495446 | B.1.1 (GR) |
| Kafkas-SARSCoV2-037 | MT787506 | hCoV-19/Turkey/KU-037/2020 | EPI_ISL_495447 | B.1.1 (GR) |
| Kafkas-SARSCoV2-038 | MT789691 | hCoV-19/Turkey/KU-038/2020 | EPI_ISL_495448 | B.1.9 (GH) |
| Kafkas-SARSCoV2-039 | MT787744 | hCoV-19/Turkey/KU-039/2020 | EPI_ISL_495449 | B.1.1 (GR) |
| Kafkas-SARSCoV2-040 | MT787746 | hCoV-19/Turkey/KU-040/2020 | EPI_ISL_495450 | B.1.1 (GR) |
| Kafkas-SARSCoV2-041 | MT789690 | hCoV-19/Turkey/KU-041/2020 | EPI_ISL_495451 | B.1 (GH) |
| Kafkas-SARSCoV2-042 | MT787503 | hCoV-19/Turkey/KU-042/2020 | EPI_ISL_495452 | B.1.1 (GR) |
| Kafkas-SARSCoV2-043 | MT789688 | hCoV-19/Turkey/KU-043/2020 | EPI_ISL_495453 | B.1.1 (GR) |
| Kafkas-SARSCoV2-044 | MT789692 | hCoV-19/Turkey/KU-044/2020 | EPI_ISL_495454 | B.1.1 (GR) |
| Kafkas-SARSCoV2-045 | MT787558 | hCoV-19/Turkey/KU-045/2020 | EPI_ISL_495455 | B.1.9 (GH) |
| Kafkas-SARSCoV2-046 | MT789694 | hCoV-19/Turkey/KU-046/2020 | EPI_ISL_495456 | B.1.1 (GR) |
| Kafkas-SARSCoV2-047 | MT787487 | hCoV-19/Turkey/KU-047/2020 | EPI_ISL_495457 | B.1.9 (GH) |

Five hundred forty-nine (549) total and 53 unique variants were detected as shown in Table 2. All 47 genomes exhibited different kinds of variants. The distinct variants consist of 274 missense, 225 synonymous, and 50 non-coding alleles (Table 2). The most common detected variants were c.1-25G>T (5’UTR), c.2772C>T (ORF1ab), c.14143C>T (ORF1ab), c.1841A>G (S) in all our genomes (n=47), c.608G>A (N), c.609G>A (N), c.610G>C (N) in 28 samples, and c.2700G>T (S), c.310G>T (ORF7a) in 23 samples. Additionally, 31 variants were detected only once. Two hundred ninety-one (53%) variants were detected in ORF1ab, which is the longest ORF consisting approximately 70% of the whole genome. The ORF1ab is cleaved into 16 nonstructural proteins (nsp). Among those, nsp3 and nsp2 had more variants in our study (n=100 and n=27, respectively. On the other hand, all non-coding mutations were detected in 5’UTR region. All genomes had c.1-25C>T nucleotide variation; however only two genomes had additional non-coding mutations which were c.1-56G>T and c.1-77G>T. In terms of base changing, the most common detected (96%) was C>T. The detailed data about coding and noncoding mutations in our isolates is in Table 2.

**Table 2:**
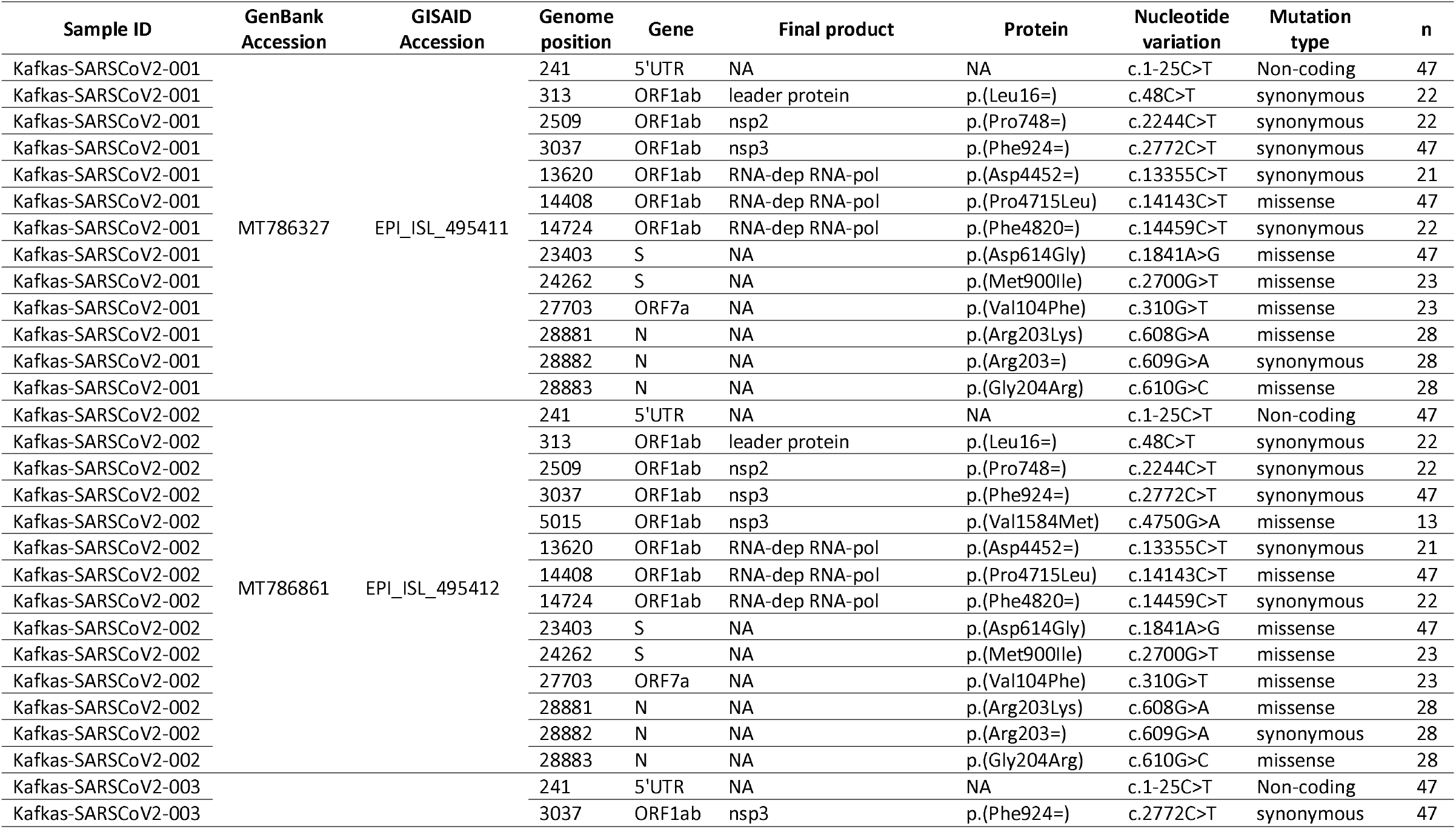
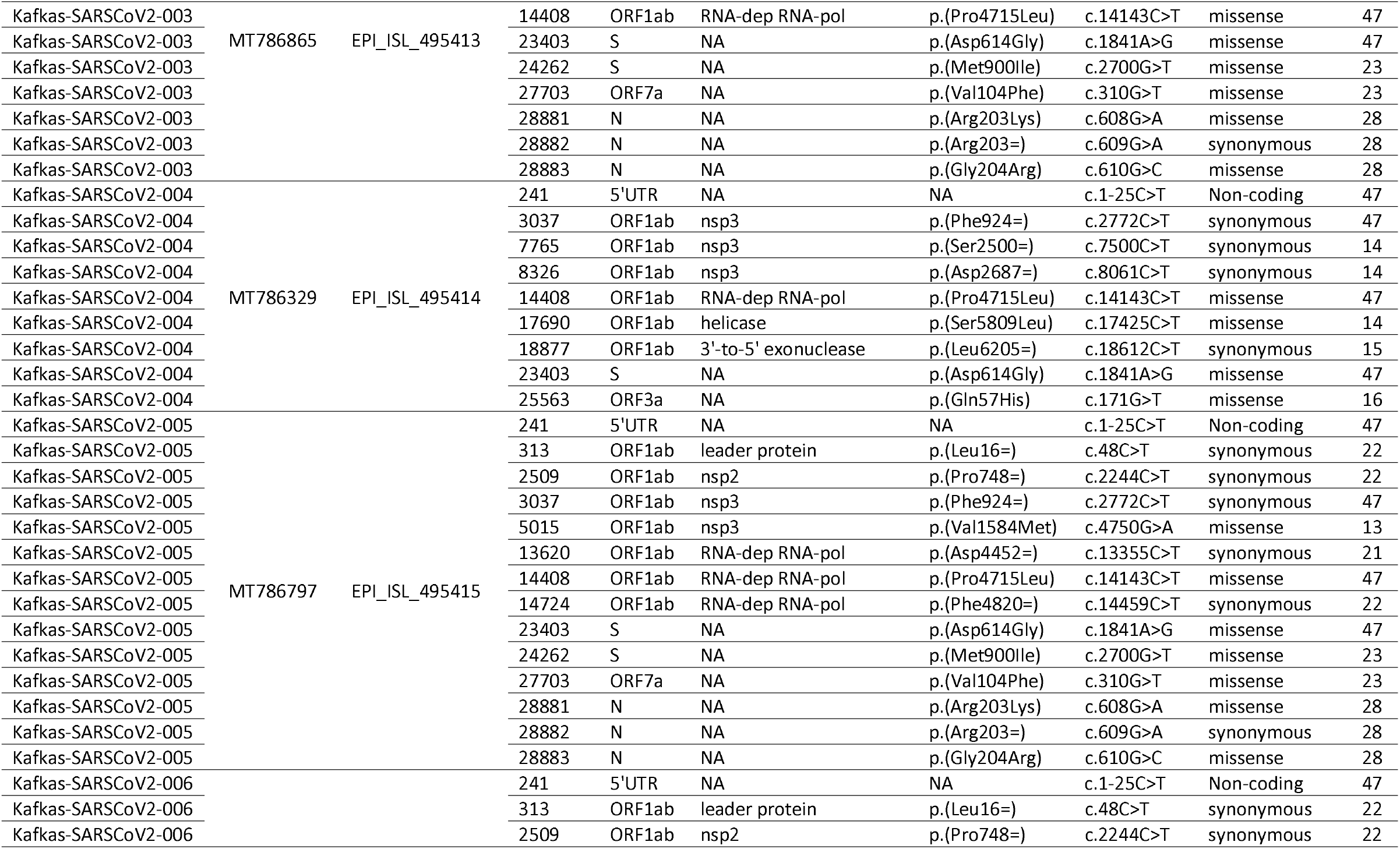
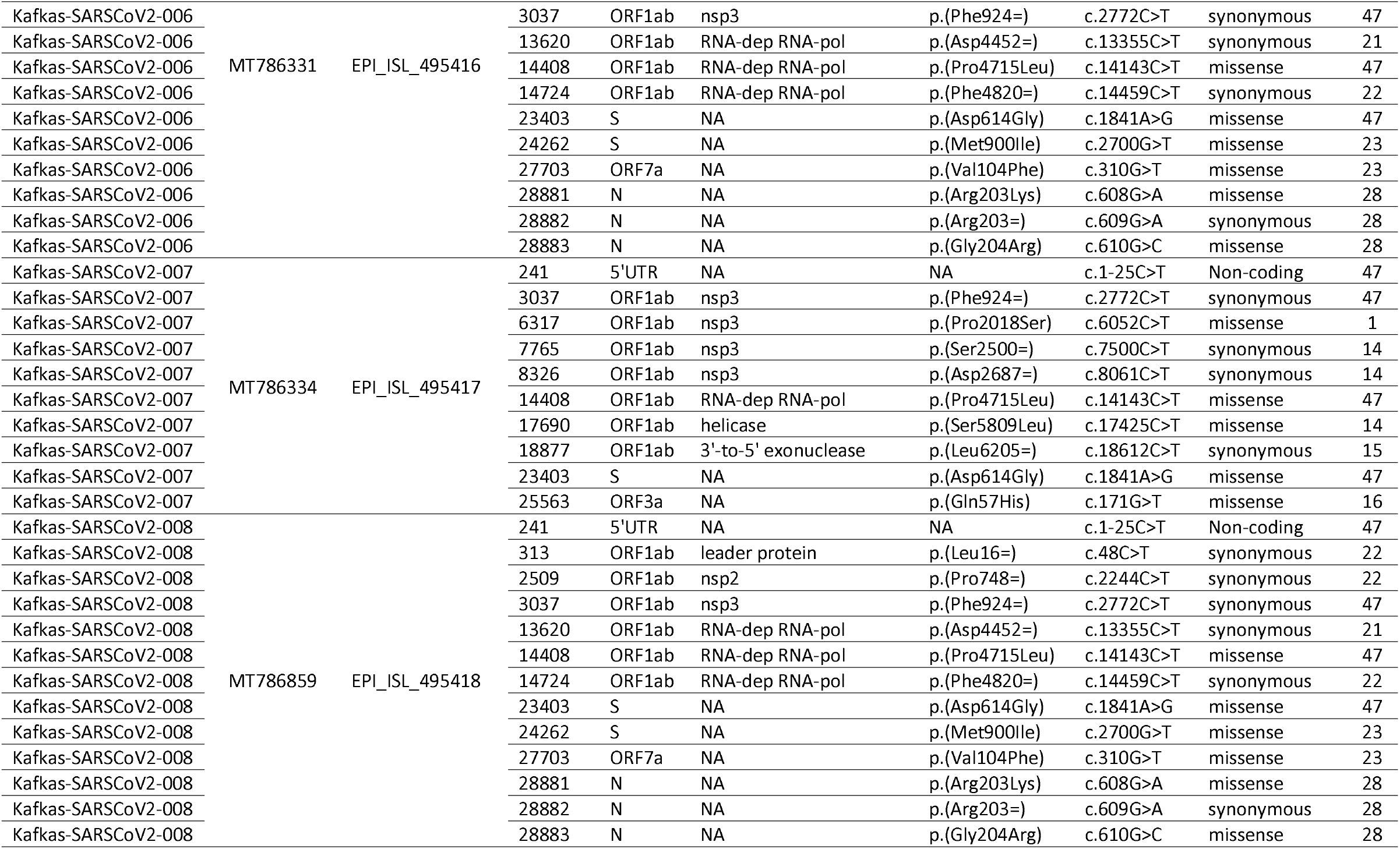
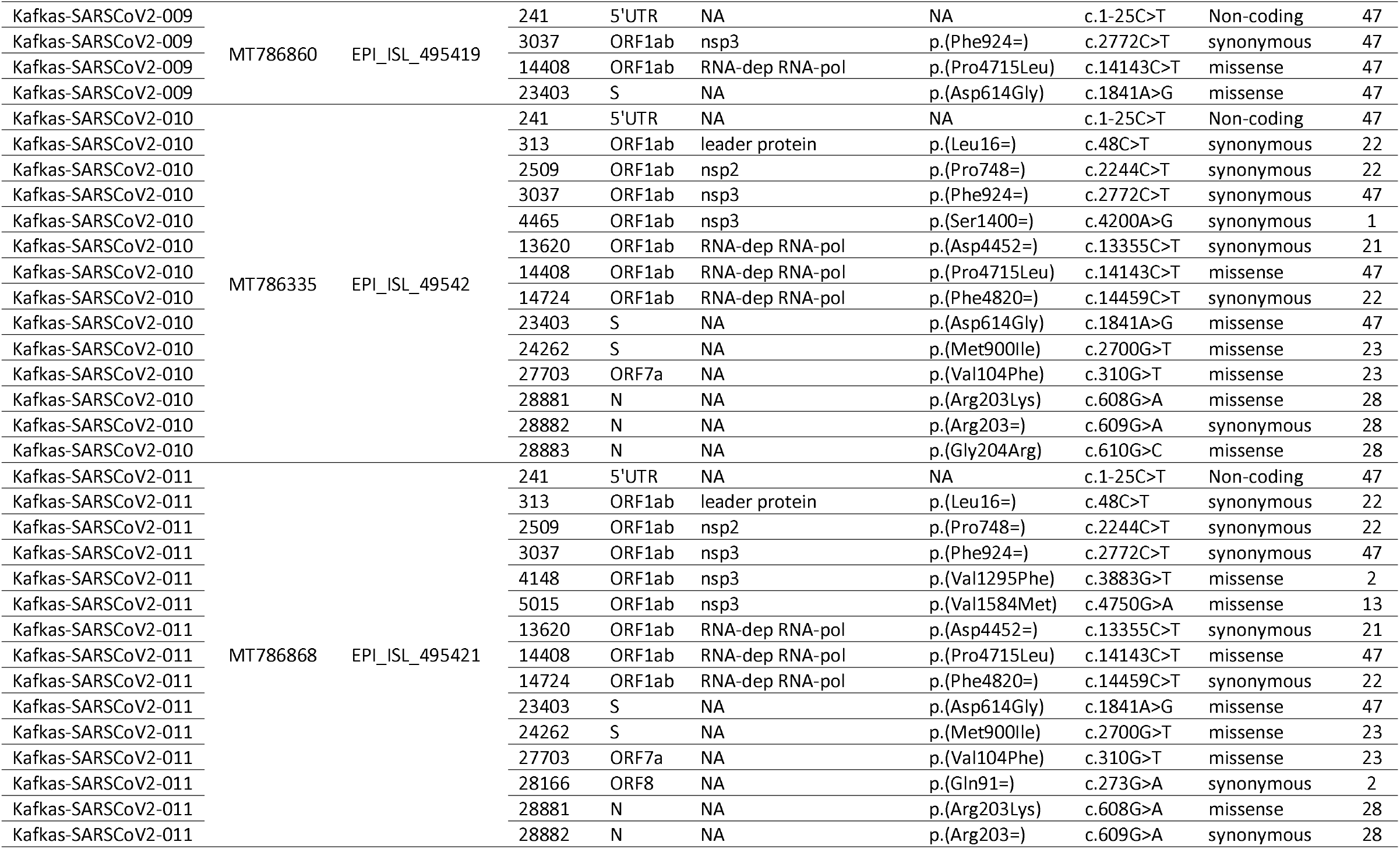
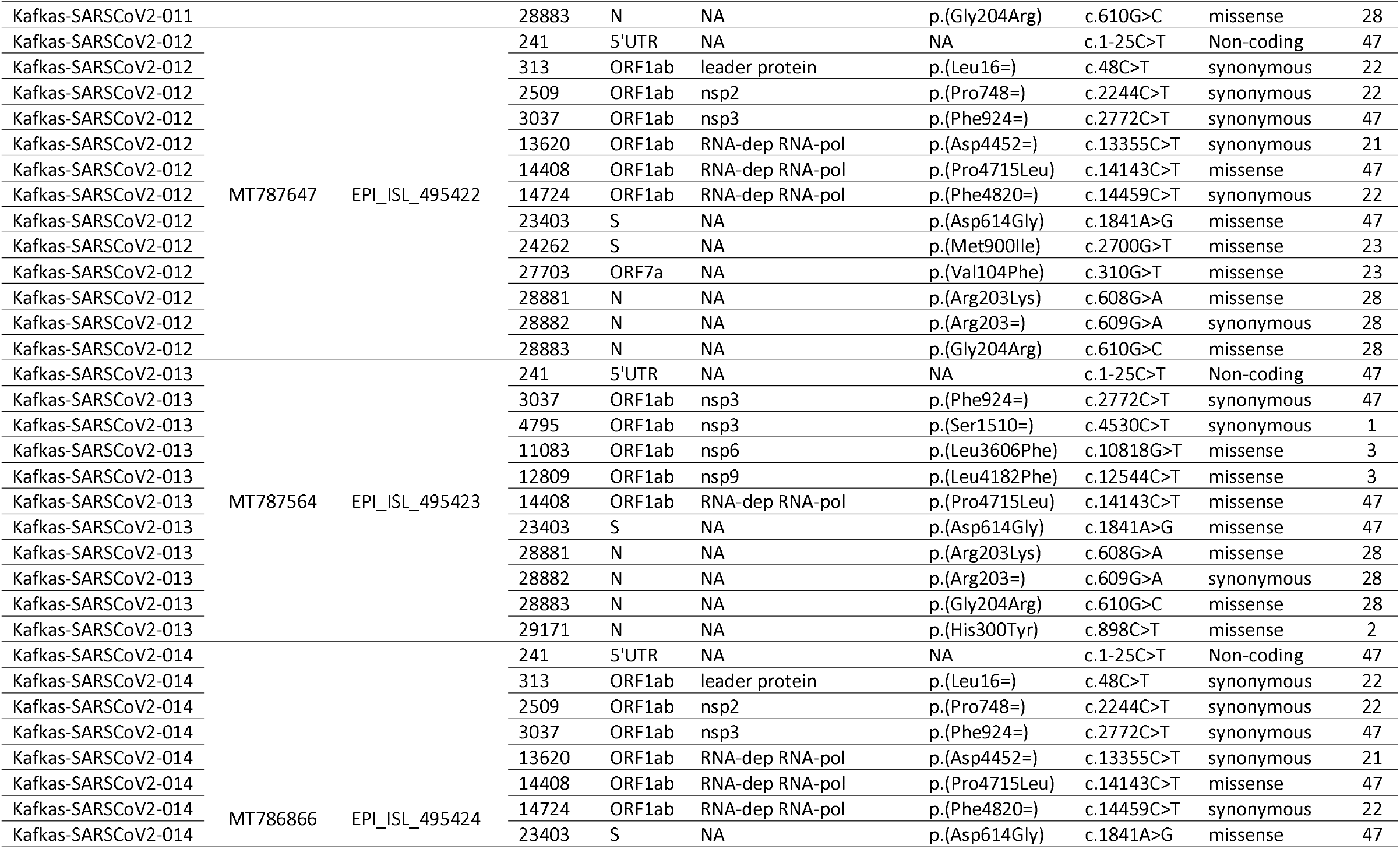
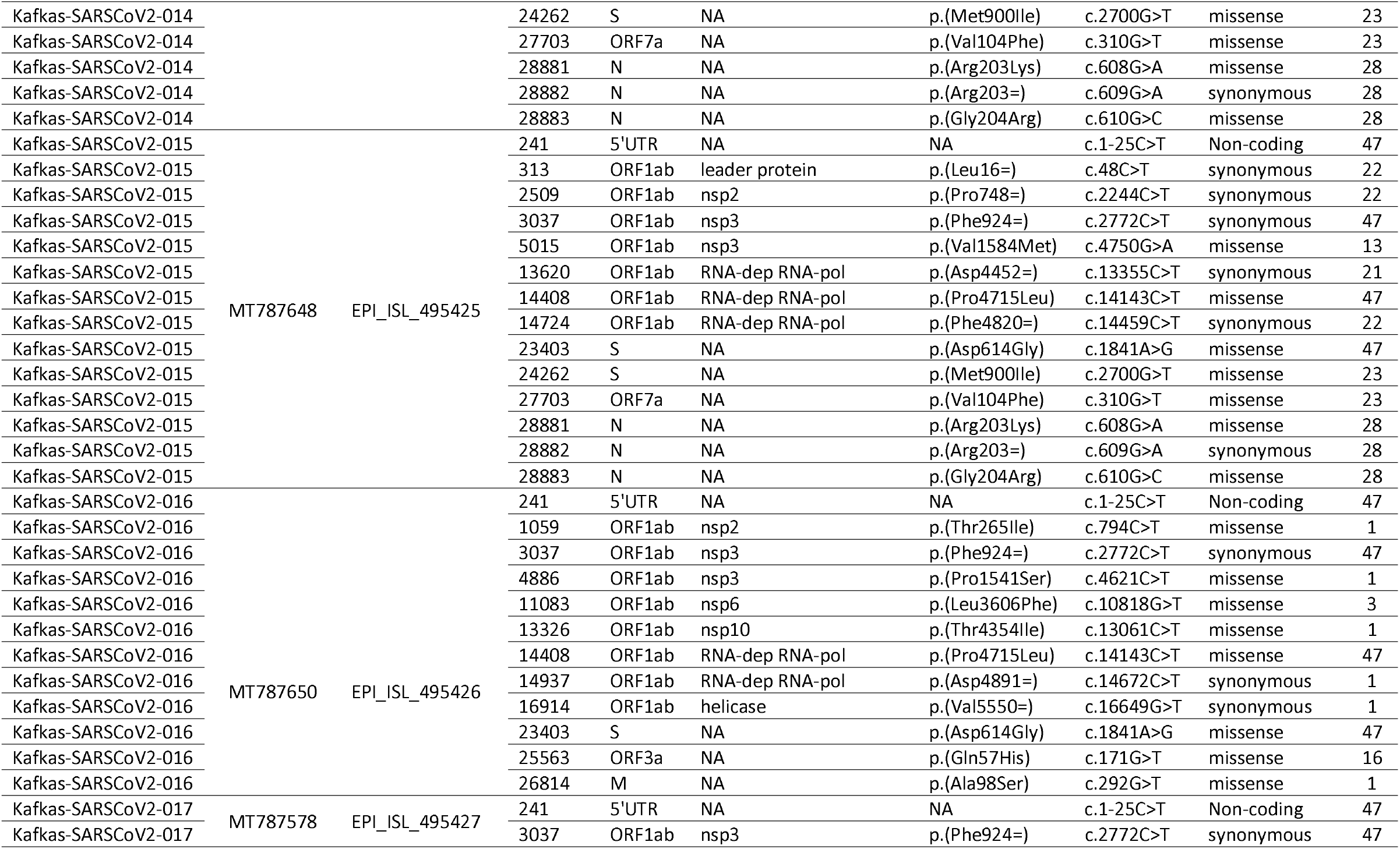
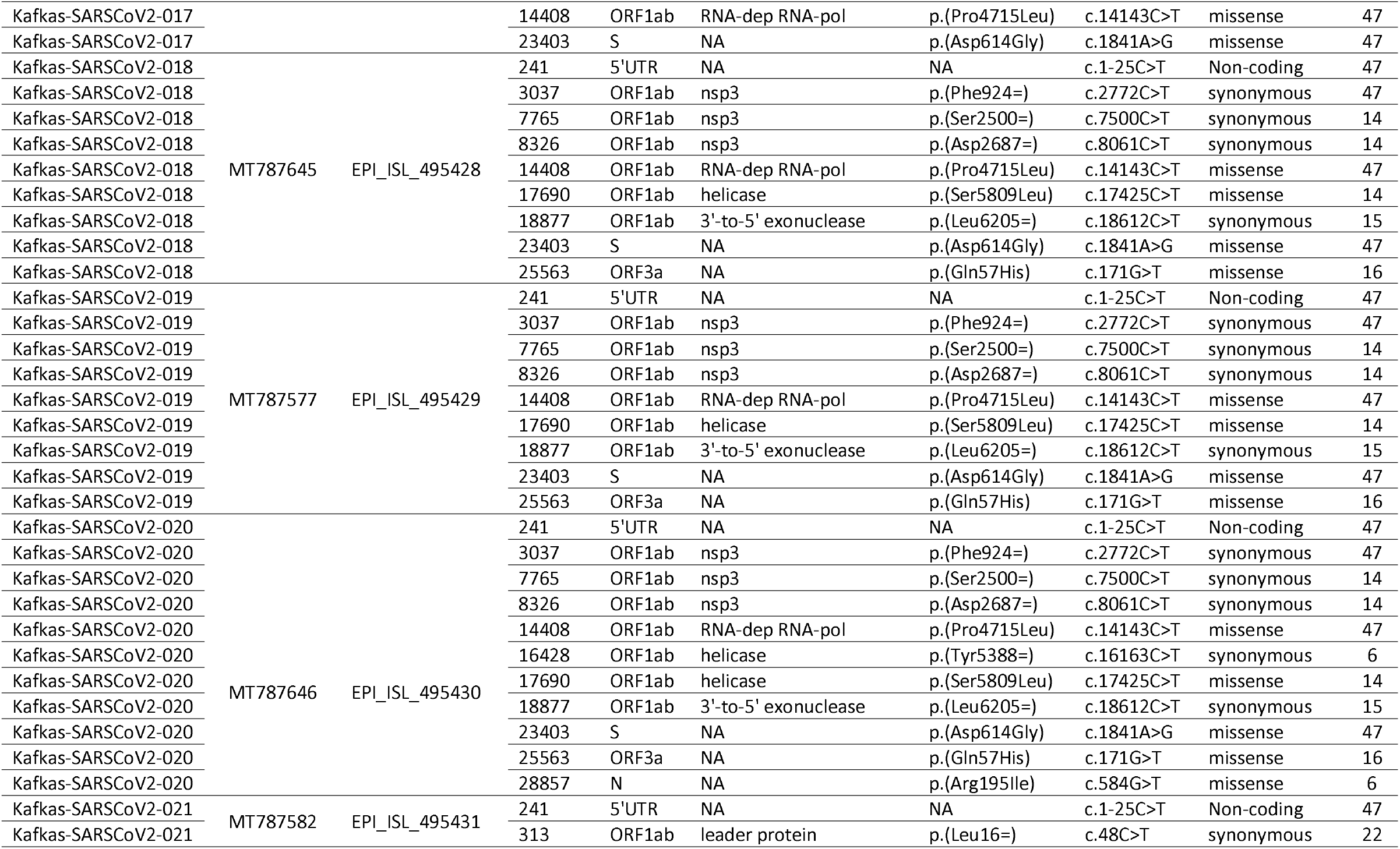
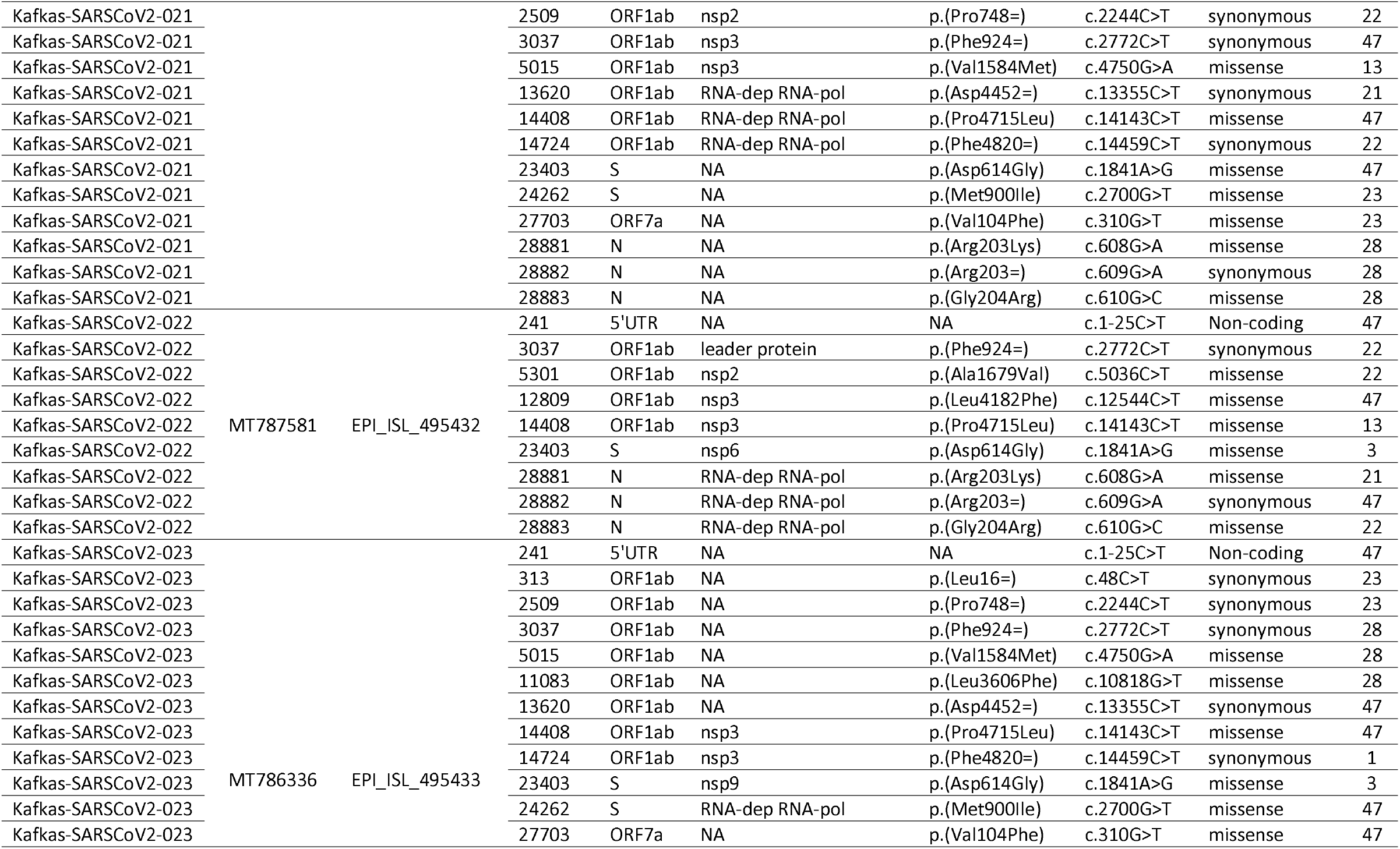
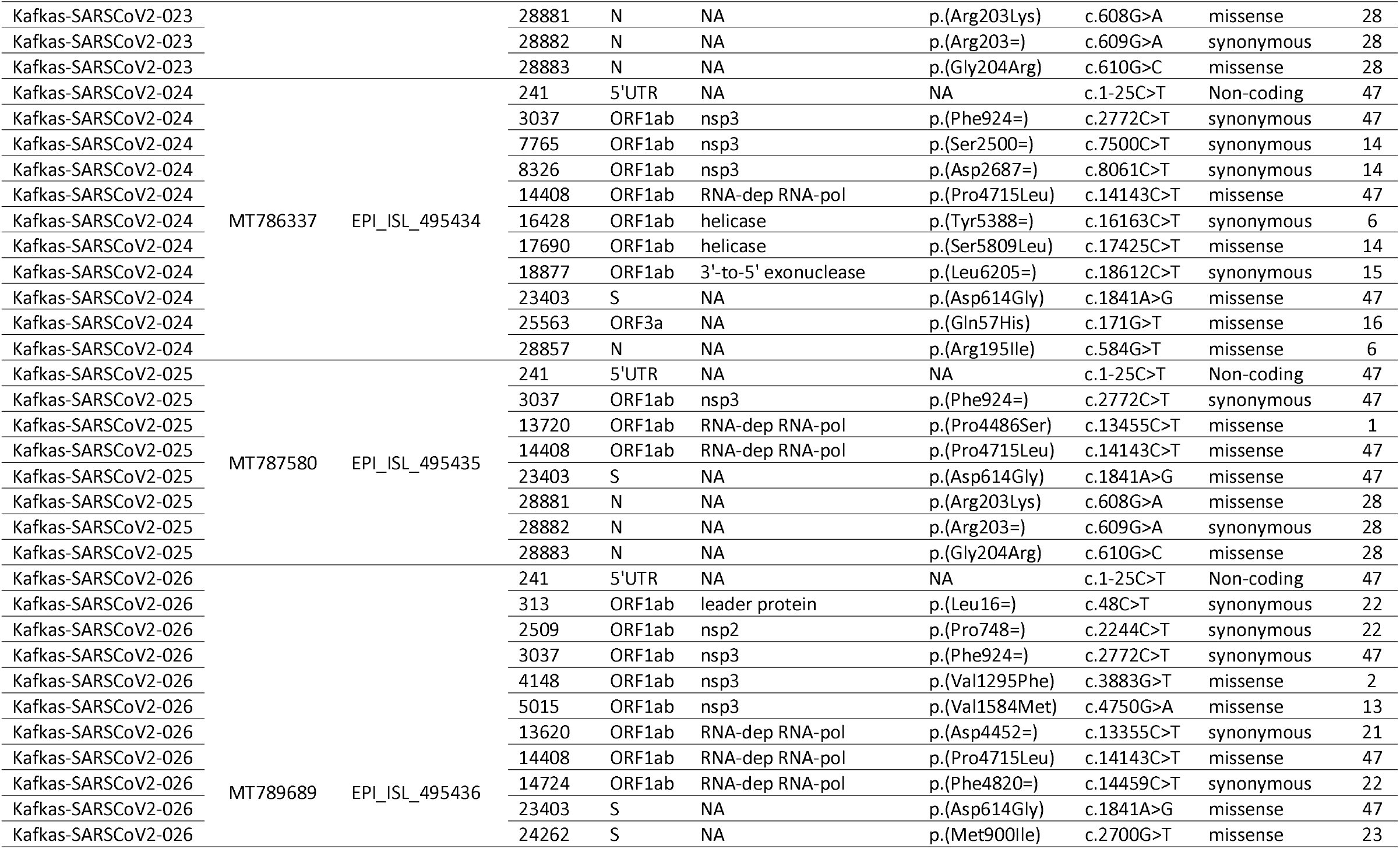
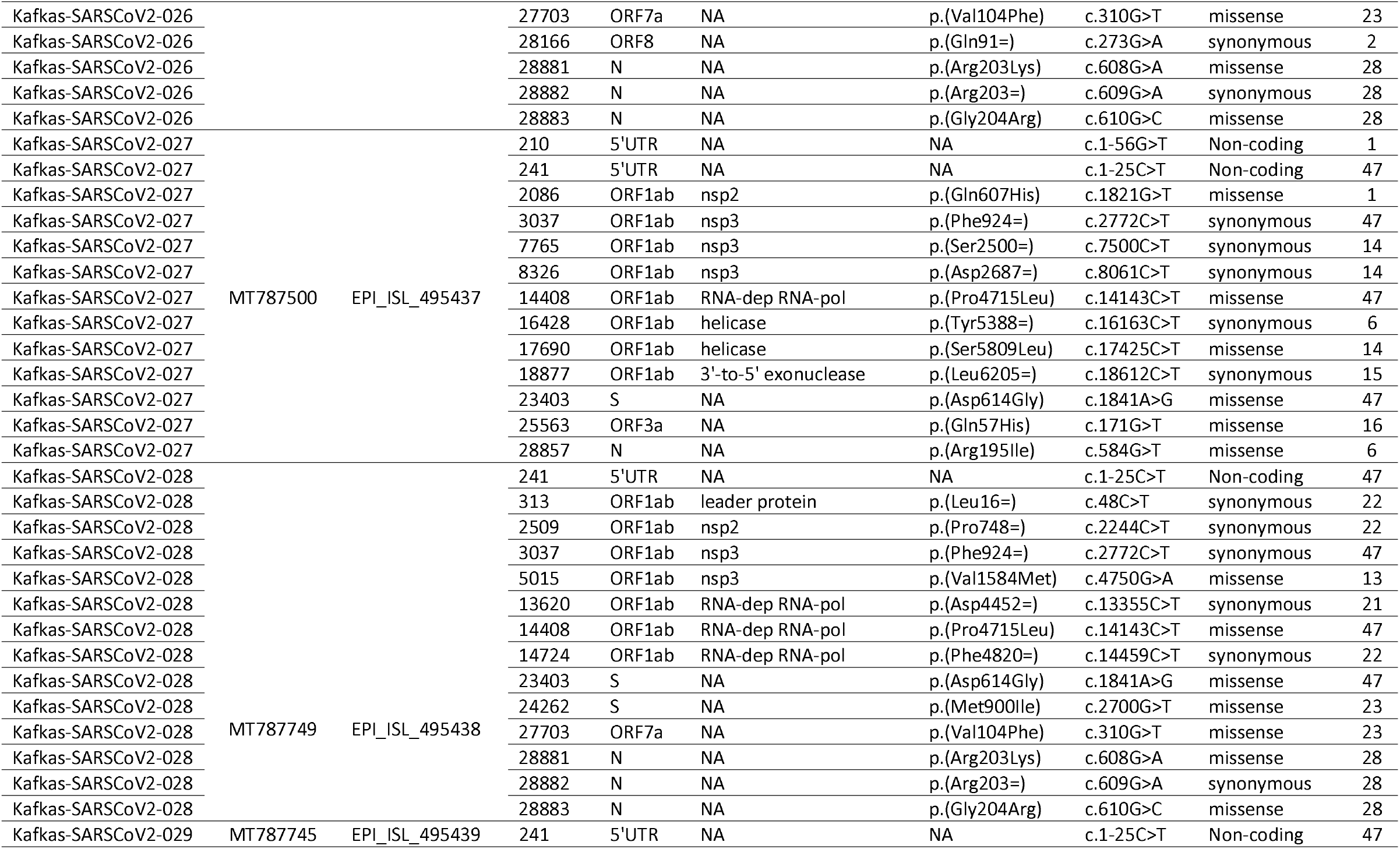
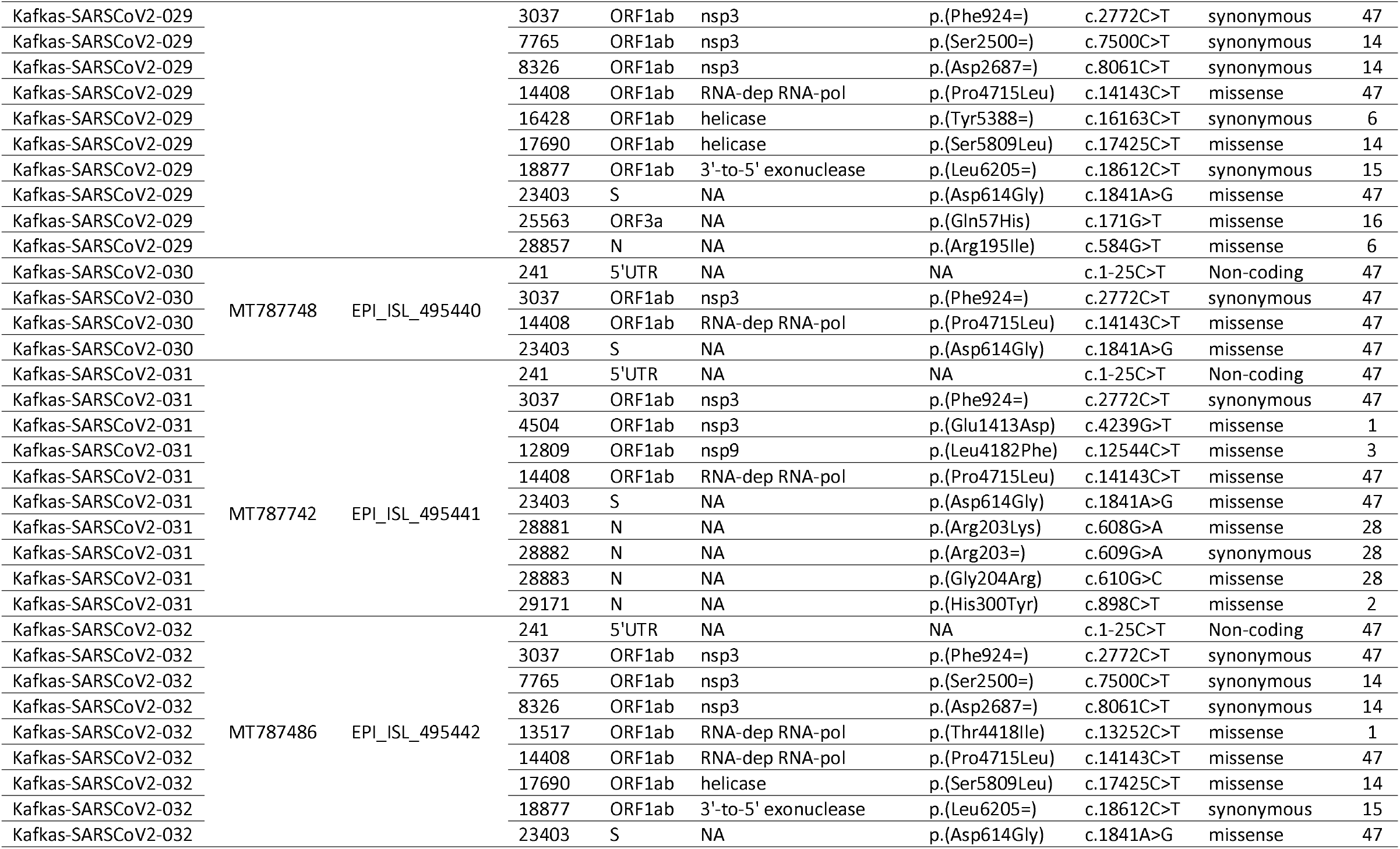
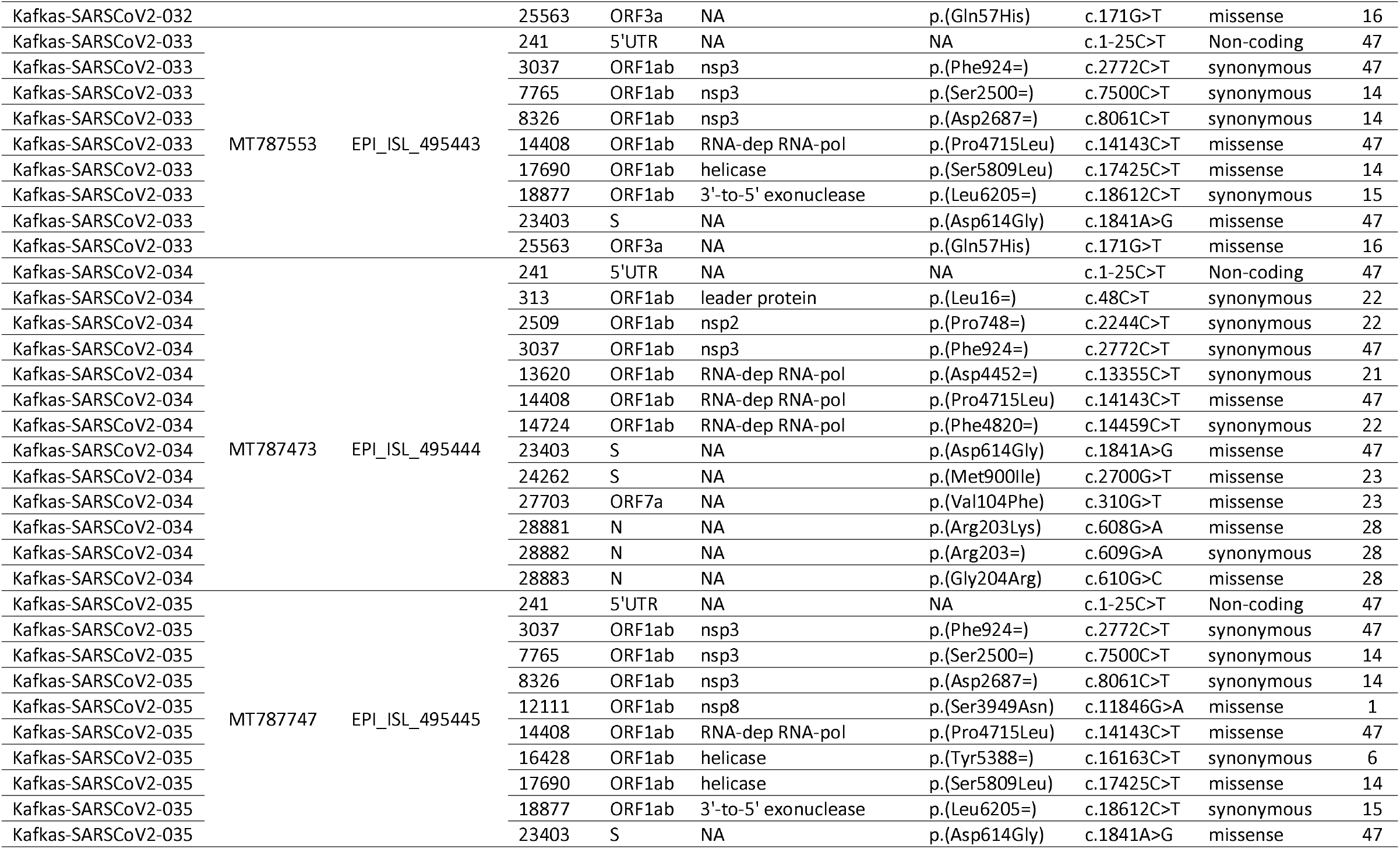
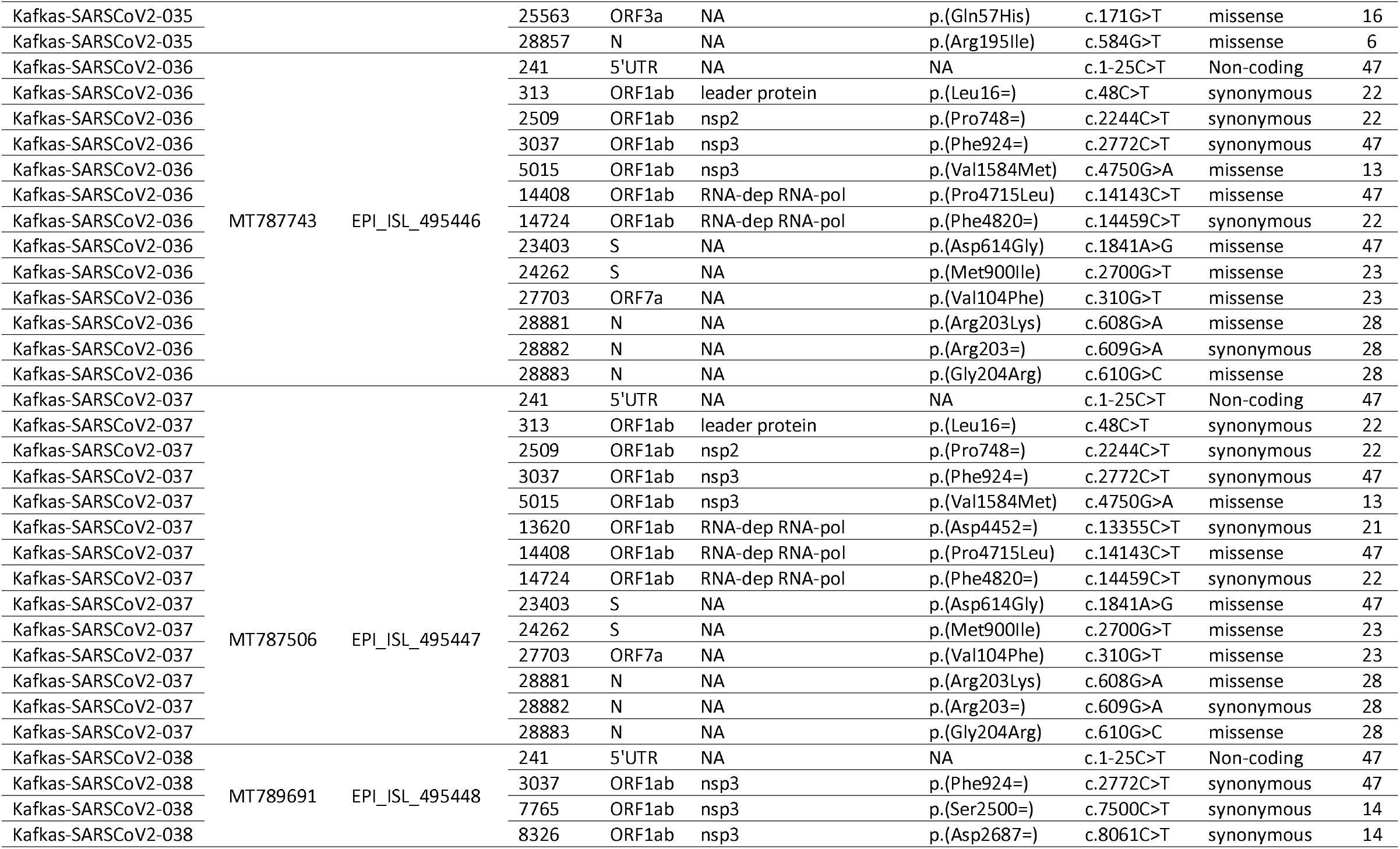
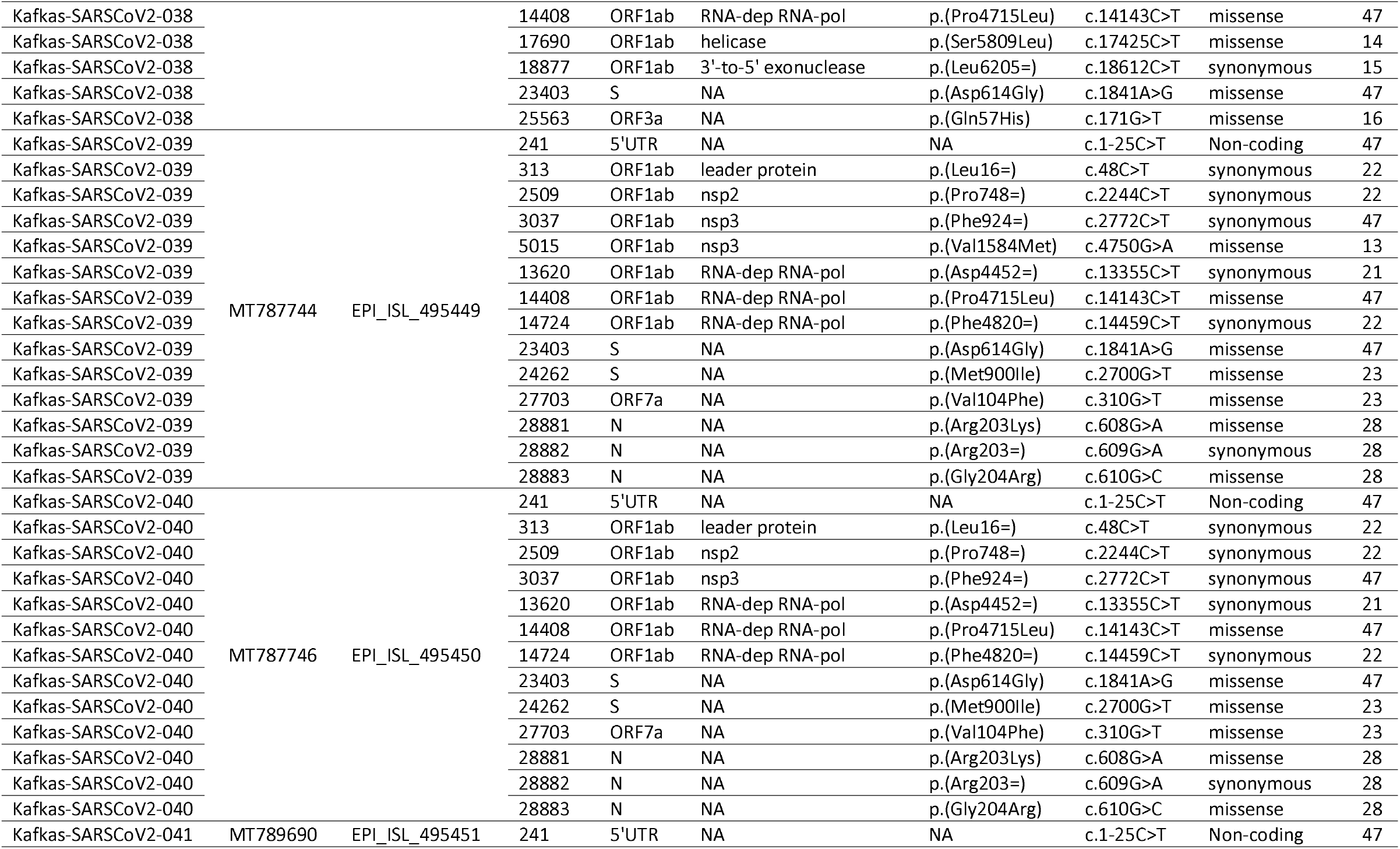
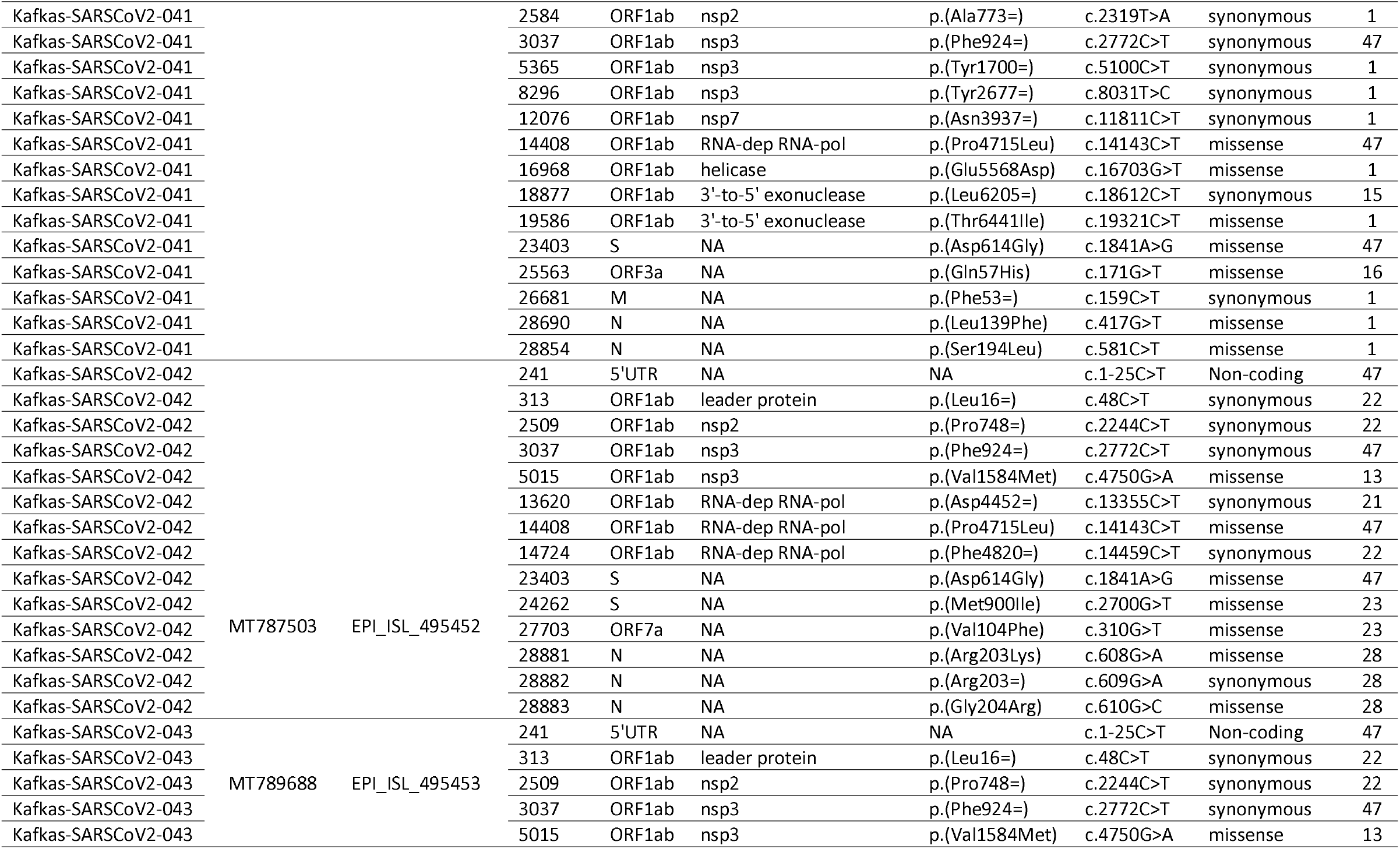
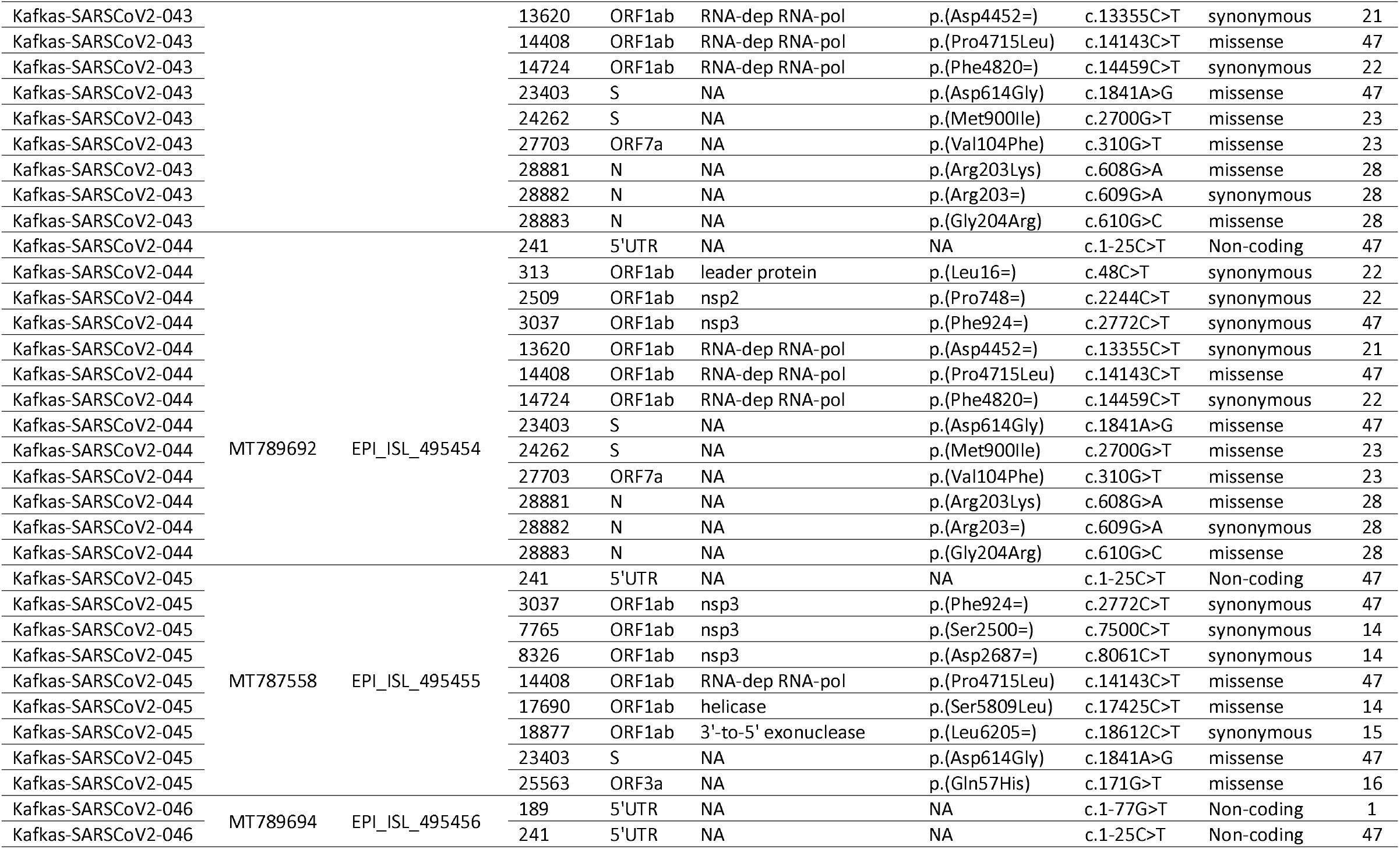
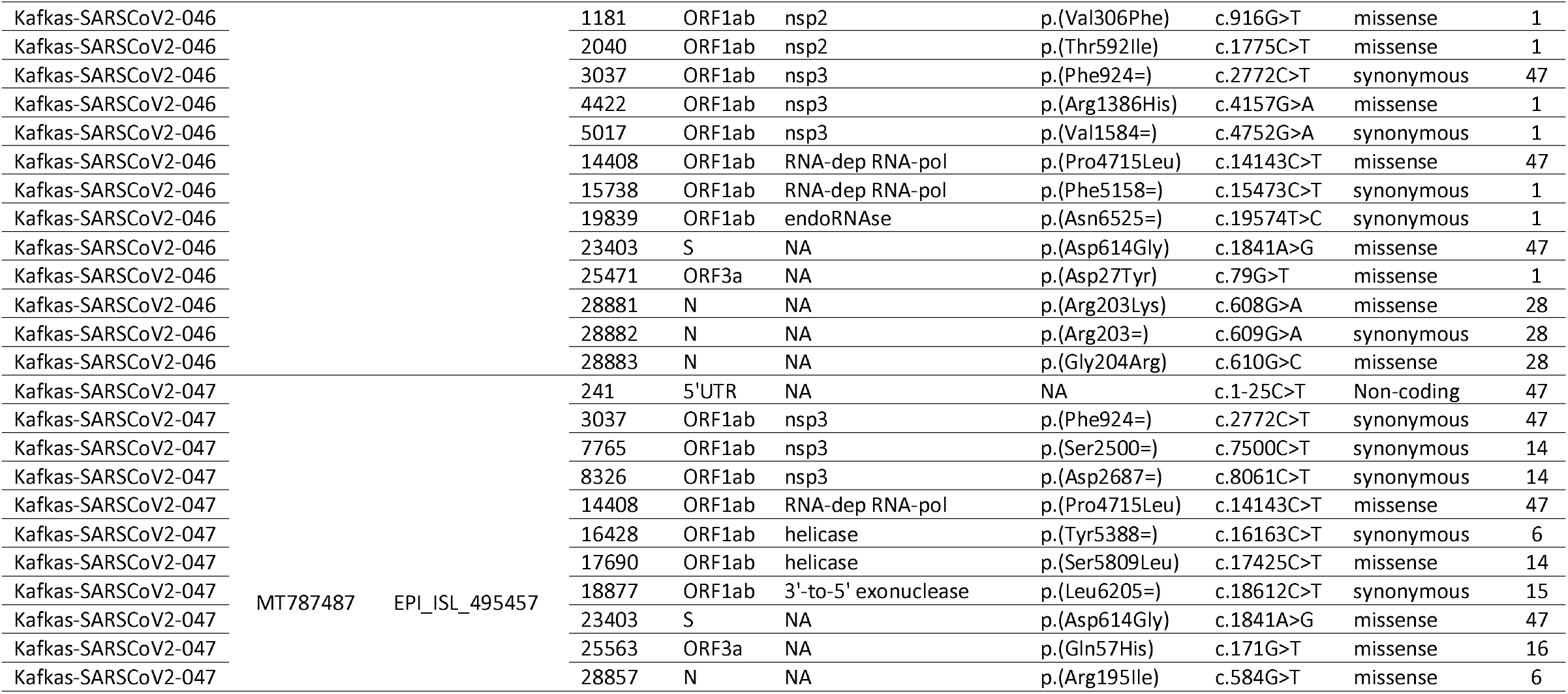
Coding and non-coding mutation list detected in SARS-CoV-2 genomes of our isolates.

## Discussion

The analysis of complete genome sequencing of new viruses is an important tool for epidemiology of infectious diseases, updating diagnostics and assessing viral evolution (7, 8). Complete genome data of a virus makes visible some epidemiological parameters including doubling time of an outbreak, reconstruction of transmission routes and the identification of possible sources and animal reservoirs. It can also help the study on drug and vaccine design. In our study, we described 47 SARS-CoV-2 genomes and compared them with a dataset of 68 available SARS-CoV-2 genomes from different countries obtained from GISAID (9).

After the phylogenetic analysis, it has been detected that all our SARS-CoV-2 genomes are similar with the European strains. According to GISAID lineage, 47 in 47 of our SARS-CoV-2 genomes are in lineage B that mostly consists of European strains. According to the marker mutations, those were placed in G, GR and GH clades. Moreover, most of European isolates have been placed in G, GR and GH clades in GISAID (9).

In current literature, there are some studies that investigate the first isolates of their country to find the originate of those (3, 11–15). A study contains two complete genomes of a Chinese patient visiting Rome and an Italian patient reported that Italian patient’s sequence clustered with European sequences and Chinese patient’s sequence clustered with Wuhan sequence (NC_045512) (16). The same researchers compared the two genomes regarding concurrent evolution and accumulation of mutations and reported that four mutations were detected in Italian sequence. Another study reported that because of the variations the first isolates of India were not showed high identity with Wuhan sequence (NC_045512) (15). Moreover, Bal et al. reported that the first three French sequences were located in European clade instead of reference Wuhan clade because of the three-nucleotide deletion in Orf1a at positions 1607-1609 (11). On the other hand, some European isolates that are generally the first isolates of related country are belonging to non-European clades. A German sequence was evaluated in a study performed by Zehender et al. (17) and reported that it belonged to lineage A because of a travelling to Shanghai between 20-24 January.

So far, there are 138 SARS-Cov-2 complete genome sequence data except our 47 isolates in GISAID database from Turkey (9). It can be seen that all those genomes are belonging to lineage B including the first case (lineage B.4) in Turkey. Our all sequences (n=47) are belonging to lineage B as similar with all other Turkish sequences. However, the first cases in Turkey is belonging to L clade according to GISAID classification whereas ours are belonging to G, GR and GH clades.

On the other hand, the length of SARS-CoV-2 genome is approximately 30 Kb including 5’UTR and 3’UTR non-coding sequence, Orf1ab, S, Orf3a, E, M, Orf6a, Orf7a, Orf7b, Orf8, N, and Orf10 genes. In this study, we aimed to detect the variations of our SARS-CoV-2 isolates.

According to the current literature, ORF genes have crucial role during COVID-19 (18). So in our study, 549 total and 53 unique variants were detected from 47 SARS-CoV-2 isolates (Table X). A study performed by Khailany et al. (19) reported that 156 total and 116 unique variants were found in 95 SARS-CoV-2 isolates. The same study also pointed out that the most frequently observed base changes was C>T same as our study.

The most observed variations are found in position 3036, 11083 and 13402 belonging to Orf1ab gene; 28854 belonging to N gene; 21707 and 21575 belonging to S gene; 28077 and 28144 belonging to Orf8 gene (4, 17, 20, 21). Our variations in related positions are same with those. We mostly detected variations in position 3037 and 11083 in Orf1ab gene; 28166 in Orf8 gene; 28854 in N gene; and 24262 in S gene. According to the global frequency of variations, most of our variants are novel; however, the studies on this issue in ongoing and more detailed information will be presented in near future. The Orf1ab is the longest and most important gene among coronaviruses (18). Most of the scientist detected the variations in Orf1ab gene. Thus, the mutations in this region may be significant concerning clinical features.

Our study have some limitations; (I) the number of national genomes available at the time of analysis; and (II) the number of detected variations. These limitations should be eliminated for further studies.

In conclusion, the results of present study indicated that the SARS-CoV-2 sequences of our isolates have great similarity with all Turkish and European sequences. Further studies should be performed for better comparison of strains, after more complete genome sequences will be released; however, these data may be useful to understand the dynamics of virus spread and may help the further vaccine and treatment studies.

On the other hand, the increase of SARS-CoV-2 cases all over the world is giving more genomes that may present some visibility of populace structure. This study showed the common and new variations in SARS-CoV-2 isolates. The fight against COVID-19 will last long time until the effective vaccine or drug will be developed. However, we believe that collecting and sharing any data about SARS-CoV-2 virus and COVID-19 will be effective and may help the related studies. At that point, we should carry on detecting the new variations.

## Declarations

### Availability of data and material

The data that support the findings of this study are openly available in NCBI GenBank and GISAID at https://submit.ncbi.nlm.nih.gov/subs/ and https://www.gisaid.org/.

## Acknowledgement

We would like to thank all patients, clinicians, and laboratory technicians who contributed to this study.

